# Developmental and genetic modulation of evidence integration dynamics in zebrafish sensorimotor decision-making

**DOI:** 10.64898/2026.03.01.708829

**Authors:** Roberto Garza, Ahmed El Hady, Armin Bahl

## Abstract

Animals integrate information over time and maintain persistent internal representations of cues to guide decision-making. How the underlying behavioral algorithms of individual animals depend on factors such as experience, developmental stage, or genotype remains poorly understood. Drift-diffusion models provide a powerful theoretical framework to describe and predict performance metrics across a wide range of species. The stochastic nature of these models and the typical limited throughput of most experimental designs challenge the automatic inference of latent variables. Here, we combine high-throughput behavioral assays in larval zebrafish with drift-diffusion modeling, revealing that larvae progressively develop more persistent, self-reinforcing integration dynamics during early development. This effect is reduced in fish carrying mutations in genes linked to human epilepsy and schizophrenia. Our results show that behavior-based drift-diffusion modeling can offer a scalable, automated approach to generate experimentally testable hypotheses about the algorithmic implementation of sensorimotor integration in health and disease.

**LINK TO VIDEOS:** https://cloud.uni-konstanz.de/index.php/s/qx9DSFciAey3aD7

**TEASER:** Evidence-integration dynamics in zebrafish mature during development and are selectively altered by disease-associated mutations.

## INTRODUCTION

In the natural world, sensory information is often dynamic and only sparsely distributed, making it challenging for animals to extract relevant cues during decision-making. A key strategy is to temporally integrate signals and thereby cancel out environmental, early perceptual, and downstream processing noise. For example, rats accumulate spatially distributed auditory cues over time to decide about the location of a water reward (*1, 2*). Mice decide to escape toward the shelter after assessing the level of threat of looming predator-like cues over time (*3*). Flies initiate a landing response after integrating repeatedly expanding low contrast patterns (*4*) and can improve odor discrimination accuracy when temporarily sampling olfactory information (*5*). Primates report their perception of overall directional movement when presented with clouds of random-dot-motion stimuli, where only a small fraction of small dots move coherently in one direction while the rest get stochastically redrawn on the screen (*6, 7*). Together, these behavioral phenomena point towards common algorithmic strategies for information processing across species.

Bounded evidence accumulation—integrating sensory cues over time until a threshold is reached—has emerged as a core computational principle underlying decision-making (*7*–*10*), describing dynamical processes across various behavioral contexts with and without reward contingencies (*11*–*14*). In this drift-diffusion framework, several model parameters, i.e., diffusion, integration leak, and evidence-based drift, control the dynamics of the decision variable and the initiation and termination of decision events. Experimentally, it is often only possible to observe the final decision, not the dynamics of the integration process itself. Thus, these underlying variables are usually hidden (latent) and must be inferred indirectly from behavioral observations. Computational modeling and reliable fitting methods make it possible to infer how these latent variables are distributed across individuals (*15*) and how they depend on developmental stage, experience, genotype, or treatment. Such insights can then help link the behavioral algorithms guiding decision-making to their neurobiological representation (*10*).

Drift-diffusion models are widely used to quantify how sensory evidence is accumulated over time during decision-making (*2, 8, 16*–*19*) and can effectively predict key behavioral features, such as decision accuracy and response delay (*20, 21, 17, 18, 9*). While these models successfully describe the behavioral algorithms, it remains debated whether model parameters map directly onto biological mechanisms (*17, 22*–*24*). Moreover, it is unclear how algorithms vary across individuals and which processes modulate parameters. The stochastic nature of these models makes it challenging to estimate underlying parameters automatically (*16*), especially when only limited trial numbers are available (*25*–*27*). Thus, previous efforts have largely resorted to manually tuning parameters (*17, 28*) or to leveraging large amounts of data for parameter inference (*2, 29*–*31*). Multi-objective fitting approaches (*32*) had some success but often produced non-unique parameter estimates across runs, complicating interpretations (*18*). To address this problem, we need experimental systems that combine high-throughput behavioral measurements and mechanistic access, supported by new parameter inference methods (*16*).

Larval zebrafish (*Danio rerio*) offer an excellent model system for studying decision-making in the context of sensorimotor processing. Not only is high-throughput single-animal tracking feasible in these vertebrates, but they also exhibit rich, quantifiable behaviors (*33*) and are amenable to molecular genetic manipulation methods (*34, 35*). A well-studied behavior in larval zebrafish is the optomotor response—the tendency of animals to align swimming with global visual motion in the environment. This behavior is thought to be an important reflex to counteract involuntary drifts in turbulent or flowing water (*36, 37*). While usually stimulated through moving gratings, it has recently been shown that the optomotor response in larval zebrafish can also be induced by random-dot-motion cues (*17, 28, 38, 39*), similar to the ones classically used in many primate perceptual decision-making studies (*6, 40*). Compared to gratings, these stimuli have the advantage that one can adjust the difficulty of the directional estimation task by varying the level of motion coherence, requiring animals to temporally integrate motion information to make directed swimming decisions (*17, 38*).

In this study, we combine drift-diffusion modeling with a Bayesian-optimization-based pipeline to automatically uncover the latent sensorimotor decision-making dynamics during random-dot-motion integration. We initially validate our method on artificially generated datasets to assess the repeatability of the parameter identification and its robustness to noise and to quantify the minimum data requirements. We then highlight inter-individual variability and link specific parameters to the developmental stage. We find that the integrator displays self-reinforcing persistence and priming-like properties when animals grow older during early development. We also probe behavior in animals carrying specific disease-related genetic defects. In larvae with mutations in *scn1lab*, a gene related to neural excitability (*41*) in human Dravet syndrome, and *disc1*, a gene encoding a schizophrenia-linked scaffolding protein (*18, 42*), the priming-like features of the integrator are largely abolished.

## RESULTS

### Zebrafish larvae show high inter-animal variability during sensorimotor integration

The drift-diffusion framework has been successful in describing evidence integration and decision-making across a large list of species. We therefore sought to assess to what extent it also explains the behavior of freely swimming larval zebrafish. If swimming statistics were captured well by drift-diffusion models, the high-throughput experimental pipelines available for larval zebrafish would enable automatic and reliable inference of the latent cognitive processes in individual animals.

Larval zebrafish move in discrete swimming events, each of which can be considered the outcome of a sensory accumulation and decision-making process. In between swims, they are largely quiescent. During this inter-swim period, animals integrate cues from the environment to guide the next motor action. We displayed random-dot-motion cues to freely swimming larvae at the age of 5 days post-fertilization (dpf) and used their optomotor response as a behavioral readout (**Fig. 1a**). To ensure controlled stimulation conditions, we locked the direction of the coherently moving dots to the location and orientation of animals in real-time (**Fig. 1b** and **Supplementary Movies 1–3**).

**Fig. 1.**
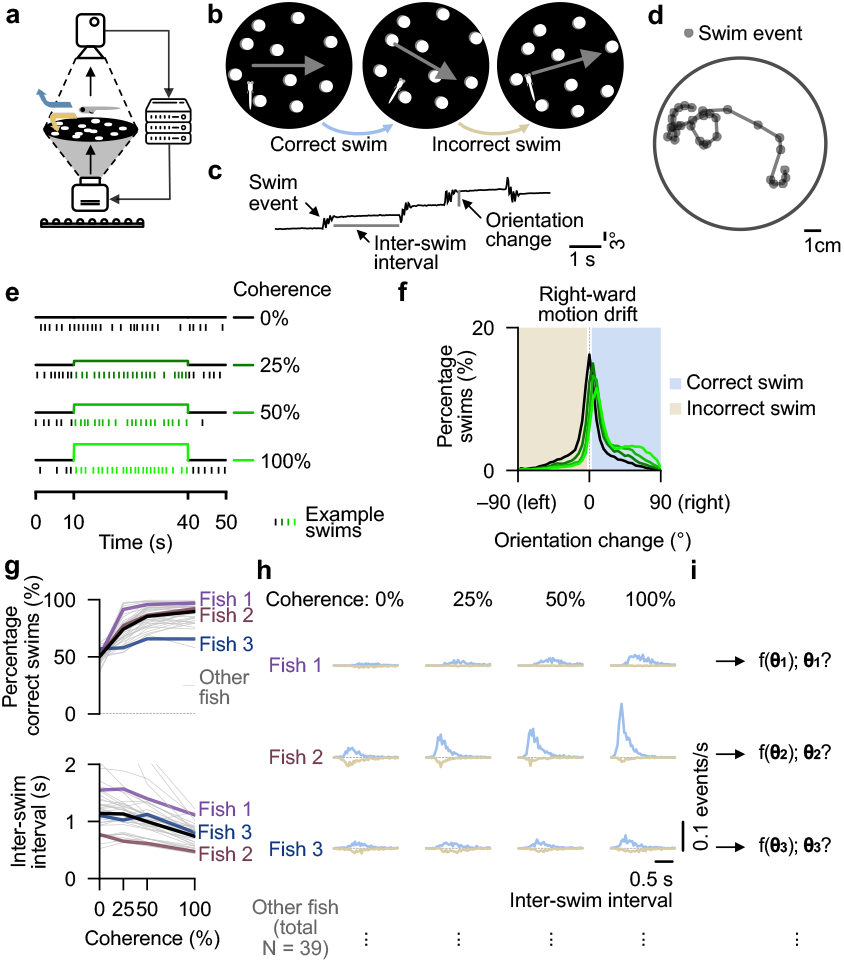
Sensorimotor integration variability suggests animal-specific latent cognitive parameters. **a**, Schematic of setup for real-time tracking of freely swimming larvae during visual stimulation. The round dish has a diameter of 12 cm and is illuminated via infrared light from below. **b**, Directed random-dot-motion at different coherence levels is presented at the bottom of the arena and is continuously locked to the orientation of the fish (**Supplementary Movies 1–3**). Motion-following turns are classified as correct swimming decisions (blue). Turns against motion as incorrect swimming decisions (gold). **c**, Example trace of animal orientation over time, used to identify and quantify individual swim events. The inter-swim interval, a proxy of response delay, is defined as the time between the end of a swim event and the start of the next one. Orientation change indicates how much animals turned during an individual swim event. **d**, Example fish trajectory. Automatically identified swim events are marked as solid gray circles. **e**, Trial structure for the tested coherence levels. The direction and coherence level are randomized for every trial. **f**, Distribution of orientation changes for different motion coherence levels in response to right-ward drifting cues. Distributions for left-ward stimuli follow the same shape but are horizontally flipped. Swim events are binarized into correct and incorrect, depending on the sign of the orientation change. Straight swims in the range of –3° to +3° are excluded from the analysis. **g**, Percentage of correct swims (top) and inter-swim interval (bottom) plotted as a function of coherence level. Thin gray lines represent individual fish. Thicker colored lines highlight three example fish. The thick black line shows the average across all animals. **h**, Distributions of inter-swim intervals for correct and incorrect swims across different coherence levels, shown for the three representative fish in (g). Distributions are normalized by the length of the experiments. The horizontal gray dashed lines indicate zero. Distributions for correct responses are shown in blue above the line, while those for incorrect responses are shown in gold facing downwards below the line. **i**, Following the idea that behavior can be captured by a general computational framework, each fish uses a set of latent model parameters, here represented by theta (θ), that our fitting pipeline seeks to uncover. Data in (f,g) from N = 39 fish.

This configuration simulates conditions in which animals would be fixed in front of a screen, a prominent configuration in classical decision-making paradigms, even though subjects are moving freely in the arena. We tracked behavior as discrete swims, quantified as orientation change and inter-swim interval (**Fig. 1c,d**), metrics that reflect decision accuracy and response delay, respectively.

We probed behavior as a function of four coherence levels, 0%, 25%, 50%, and 100%. We followed a classical stimulus design (*6, 43*), commonly used in psychophysical experiments, in which a large number of dots get repositioned at regular intervals. Here, at every frame, the coherent fraction of dots moves spatially slightly offset relative to their previous location, while the rest disappear and randomly reappear somewhere else in the arena (**Methods**). On average, swim events occurred approximately once per second during the 30-second-long stimulation period within each trial (**Fig. 1e**). We binarized orientation change into correct and incorrect swims, depending on whether animals followed motion direction or went against it, respectively (**Fig. 1f**).

Across individual animals, we found the percentage of correct swims to increase and the inter-swim interval to decrease with coherence levels (**Fig. 1g**), corroborating previously observed trends at the population level (*17, 28, 38*). To further quantify responses for individuals, we analyzed the distributions of inter-swim intervals classified as correct and incorrect as a function of coherence level (**Fig. 1h**). As expected, larvae displayed more correct swims with shorter inter-swim intervals for higher coherence levels. Quantification of the coefficients of variation (*CV*) as a function of coherence level showed that animal-to-animal variability was considerable for the average inter-swim interval but more conserved for the correctness of swims (**Extended Data Fig. 1a**). This result indicates distinct animal-specific motion integration dynamics and sensorimotor decision-making strategies and highlights the importance of analyzing and modeling behavior at the level of individual animals, rather than from a population average perspective.

We hypothesize that behavior follows a general sensorimotor integration model, however, with different individual-specific parameterizations (**Fig. 1i**). Through computational modeling, it should be possible to use the inter-swim interval distributions as target datasets to estimate latent model parameters. These extracted variables can then provide hints about specific mechanistic differences in the brain. We next developed a robust fitting algorithm to achieve these goals.

### Inferring model-derived latent cognitive variables using Bayesian optimization

The drift-diffusion model class is a simple yet prominent framework to predict key features of evidence accumulation and decision-making across various species, including primates, rodents, fish, and insects (*1, 2, 8, 17, 44, 45*). Such models have recently been shown to be generally suitable also in larval zebrafish in the context of optomotor behavior in response to temporally integrated random-dot-motion (*17, 18, 28, 38*). The previous models developed for larval zebrafish used stochastic initiation of swims with different probabilities below and above the decision bound (*17, 18, 28*). This configuration may lead to less intuitive parameter interpretations and complicate automatic parameter inference (*18*). We therefore implemented a simpler, classical (*8*) version of the drift-diffusion model with five free parameters (**Fig. 2a,b**): the diffusion coefficient (*σ*), summarizing the overall noise in the integration process; the drift rate (*µ*), which reflects the weight of sensory input; the integrator leak (λ), which determines how previously accumulated evidence alters the integration dynamics; the reset factor (*r*), governing the integrator variable state after a decision, and, finally, the delay (*δ*), accounting for the sensory or motor refractory period following a decision. The decision boundary (*B*) was held constant at *B* = 1 for all simulations, as this is a mathematically redundant parameter that scales with other integration parameters (**Methods**).

**Fig. 2.**
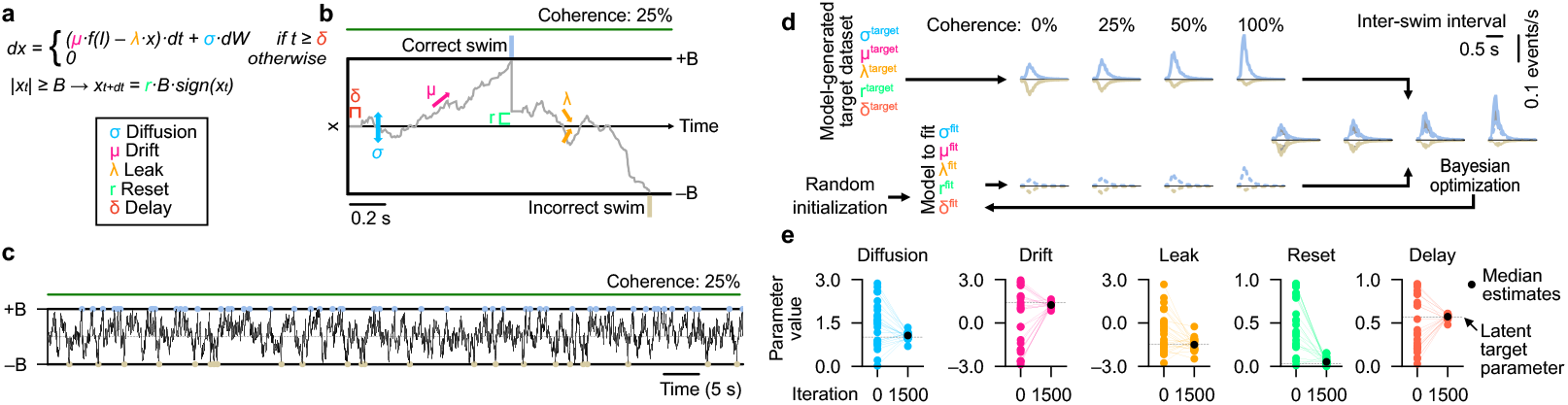
Reliable recovery of latent drift-diffusion parameters from model-generated target data. **a**, Model differential equation for the integrator variable (*x*) with color-highlighted parameters: Diffusion (*σ*) in blue, drift (*µ*) in magenta, leak (λ) in yellow, reset (*r*) in green, and delay (*δ*) in orange. Decision boundary (*B*) in black, which is a redundant parameter and kept constant at *B* = 1. *f*(*I*) describes a simple non-linear transformation of the input *I. dW* is a white noise Gaussian process, scaled with the simulation timestamp *dt*. The color code is kept consistent throughout the manuscript. See **Methods** for implementation details. **b**, Schematic representation of the model with an example trajectory of the integrator variable *x* in response to 25% motion coherence. A correct decision (blue tick) is followed by an incorrect one (golden tick). The contribution of each parameter is represented by arrows and brackets. **c**, Simulation of the model for 60 s with an exemplary selected parameter set in response to 25% motion coherence. **d**, Illustration of the fitting procedure applied to a model-generated target dataset. Solid lines are target model data. Dashed lines represent the to-be-fitted model. **e**, Initial and final parameter estimations for an example model fitted repeatedly 20 times with random initializations. Thin colored lines represent optimization runs, and black dots show the median final estimation. Gray dashed horizontal lines highlight the latent parameter values used to generate the original target behavioral dataset. The optimization algorithm reliably recovers values for each parameter. See also **Extended Data Figs. 1–3**.

To probe if our model can reliably be fit to individual behavioral datasets, we first employed a strategy for retrieving parameters from artificially generated decision events. To this end, we chose model parameter sets that can generate behavior resembling the experimentally observed regime (**Methods**). Inferring these chosen, known, latent parameters from behavior alone then allows us to quantify the performance of our fitting algorithms.

Specifically, we simulated a given target model for different levels of coherence over periods similar to the ones used in the real experiments (**Fig. 2c**), allowing us to extract a list of decision events. We then computed histograms of correct and incorrect decisions, generating the target quantities for our fitting algorithm (**Fig. 2d**). We then performed model fitting via Bayesian optimization (**Methods**), minimizing the distance between test and target model results. To compute distances between histograms, we used a modified version of the Kullback-Leibler divergence (*D*_*KL*_), which we term 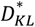 (**Methods**). This modified loss function improves the standard distance metric by weighting each bin of the histogram proportionally to its height, such that frequently occurring events are prioritized during fitting. To ensure the best results when fitting experimental data, we quantified the performance of our modified loss function using real fish. Both 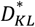 and 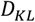 led to comparable convergence to near-zero loss (**Extended Data Fig. 1b**) but 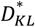 displayed better inter-swim interval matches at the end of the optimization process, in particular for low coherence levels (**Extended Data Fig. 1c,d**).

The temporal resolution of simulating the decision variable in our model largely determines the simulation time requirements of our fitting procedure, which is critical for reasonably fast optimization. We therefore validated how the length of the simulation time step affects our results. We found no benefits for values smaller than *dt* = 0.01*s*, but considerably undersampled inter-swim interval distributions for values larger than *dt* = 0.1*s* (**Extended Data Fig. 1e,f**).

After validating our optimization algorithm, we then chose an example parameterization in our model to generate artificial behavioral data. We found that the fitting procedure allowed us to reliably retrieve the chosen latent parameter set (**Fig. 2d,e**). To quantify whether parameter inference also works across the entire parameter space, we generated a library of 100 randomly generated models (**Extended Data Fig. 2a**). Each of these models could generate behavior resembling what we observed experimentally, following a list of specific inclusion criteria (**Methods**), such as that decision events happen at biologically plausible rates and that performance increases with coherence. For all tested models, we found loss functions to rapidly decay toward zero and latent parameters to be reliably retrieved, with most models having near-zero error between the target parameter set and the inferred one (**Extended Data Fig. 2b**).

We next sought to quantify how much experimental data one would need to reliably retrieve parameters (**Extended Data Fig. 2c**). The less time required to measure real animals, the higher the experimental throughput. To this end, we simulated our library of models for different lengths of time. We found that just 900 s of data per coherence (3,600 s for all four coherence levels) was enough to reliably retrieve model parameters. This value matches our experimental design (**Fig. 1e**), in which we also showed at least 900 s per coherence organized in trials with a 30-s long stimulation window (**Methods**).

Finally, we wanted to know to what extent noise in the measured inter-swim interval histograms would affect results (**Extended Data Fig. 3a**). For all experimentally realistic noise levels that we artificially added to the distributions, distance metrics converged to zero, and parameter sets could be reliably retrieved (**Extended Data Fig. 3b–h**).

To further assess the sensitivity of our estimation strategy and model robustness, we examined how perturbations of individual model parameters affected the loss function. This analysis showed particularly high sensitivity to small changes in the diffusion parameter, whereas perturbations of the other parameters had weaker effects (**Extended Data Fig. 3i**). Further statistical analysis (Mann-Whitney U test) showed that we could distinguish losses when we varied the diffusion value by ±5%. For the reset parameter, ±25% variation was necessary to generate significant effects (**Extended Data Fig. 3i**). These results indicate that our fitting algorithm navigates a near-flat region of the loss function around the optimal parameters, explaining the residual variation in parameter identifiability at the end of the optimization process (**Extended Data Fig. 2b**).

Thus, our validation approach of using model-generated data with known parameter sets clearly shows that a Bayesian fitting strategy with an improved distance metric can reliably infer latent parameters of drift-diffusion models. These results hold for experimentally feasible time scales and realistic noise levels, indicating that our method is readily applicable to behavioral data from real animals under different experimental conditions, which we explored next.

### Extraction of drift-diffusion model parameters from real experimental data

We found that our Bayesian optimization approach can reliably fit drift-diffusion models to experimental data and faithfully extract the underlying latent cognitive parameters. As a first application, we employed this validated strategy to capture the animal-to-animal variability we observed earlier (**Fig. 1g,h**) and to explore corresponding parameter variations. Compared to our model-generated approach (**Fig. 2c)**—where the underlying parameter set is known—the target data now comes from real experimental animals, for which parameters need to be unraveled (**Fig. 3a**). Procedures are otherwise identical. The extracted variables may then hint toward individual-specific cognitive mechanisms. We first tested the three previously illustrated example fish (**Fig. 1g–i**). For each animal, the loss function converged to near-zero (**Fig. 3b**), illustrating that the optimization algorithm also works for real data. The key behavioral features, measured by the percentage of correct swims and the inter-swim interval as a function of coherence level, were captured well (**Fig. 3c**). As expected from the design of the loss function and its convergence, the estimated models also matched the distributions of inter-swim intervals for correct and incorrect swims across different coherence levels (**Fig. 3d**). The success of the Bayesian optimization procedure enabled us to extract latent parameters for each of the three example fish, which showed clear fish-to-fish differences (**Fig. 3e**).

**Fig. 3.**
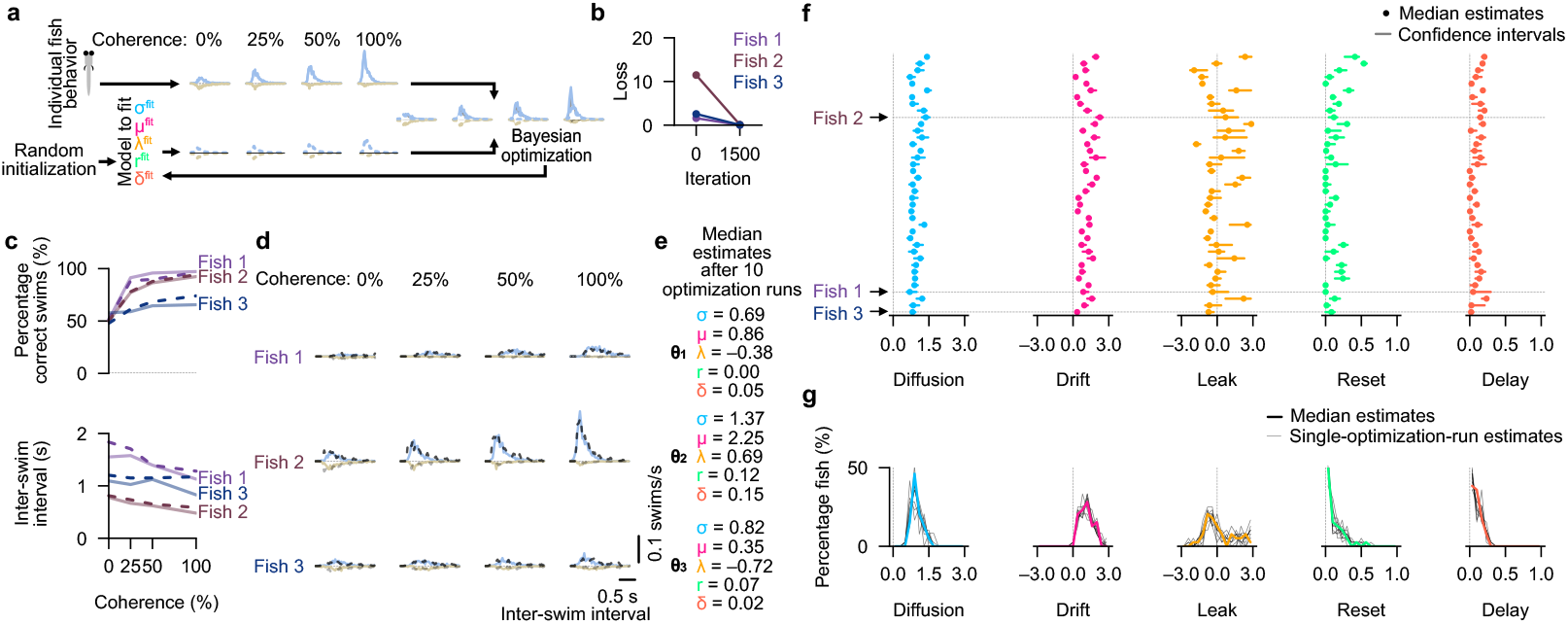
Unraveled latent cognitive variables of the drift-diffusion integration process in individuals show animal-to-animal variability. **a**, Bayesian optimization procedure for latent variable estimation, as in **Fig. 2d**, but for real zebrafish larvae. **b**, Convergence to near-zero values of the loss function between the start and end of the fitting procedure for three example fish. One optimization run per fish. Same example fish as in **Fig. 1g–i. c**, Percentage of correct swims (top) and inter-swim interval (bottom) as a function of coherence level. Optimized models reproduce the three example individuals. Experimental data are shown as solid lines, model data as dashed lines. **d**, Distributions of inter-swim intervals for correct and incorrect swims across different coherence levels for the three example fish. Solid lines indicate experimental data. Dashed lines represent fitted model results. **e**, Unraveled latent cognitive variables for the three example fish, hinting toward potential mechanistic differences across animals. Optimization was run 10 times to compute the median parameters. See confidence intervals in (f). **f**, Estimated parameter sets for all tested larvae. Each row is one individual animal (fish ID). Dashed vertical lines mark zero values. Small arrows and horizontal dashed lines indicate the three example fish displayed in (c,d). Confidence intervals describe the range between the 10^th^ and 90^th^ percentiles of the distribution originating from 10 repeated optimization runs per fish. **g**, Population estimate distributions of parameters across all tested larvae. Colored lines are the results from the medians of 10 repeated parameter estimations per fish, while black thin lines show distributions composed of single optimization runs per fish. N=39 fish in (f,g). All animals in this figure were 5 dpf old. See also **Extended Data Fig. 4**.

We then applied the strategy to all experimentally tested 5 dpf larvae. For each animal, the loss function converged to near-zero values (**Fig. 4a** and **Extended Data Fig. 1b**). Multiple optimization runs per individual led to robust repeatable estimates of model parameters with similar values (**Fig. 3f**). The extracted median parameters varied across the population (**Fig. 3f,g**). Population analysis based on single optimization runs resulted in similarly shaped distributions (**Fig. 3g)**. Thus, population-level comparative insights (see sections below) can be achieved with single optimization runs, saving computing power. Notably, in more than half of the individuals, we found the leak parameter λ to be negative. Given our formulation of the model (**Fig. 2a**), a negative leak implies that the current value of the integrator variable *x* reinforces itself, promoting persistent internal states and accelerating threshold crossings. Biologically, such dynamics are consistent with recurrent positive feedback within the accumulator, reminiscent of working-memory-like or priming-related neural processes (*2, 46, 47*). Importantly, because the integrator operates within absorbing bounds and is partially reset after each decision event, these self-reinforcing dynamics do not lead to runaway divergence but instead produce stable, bounded decision trajectories.

**Fig. 4.**
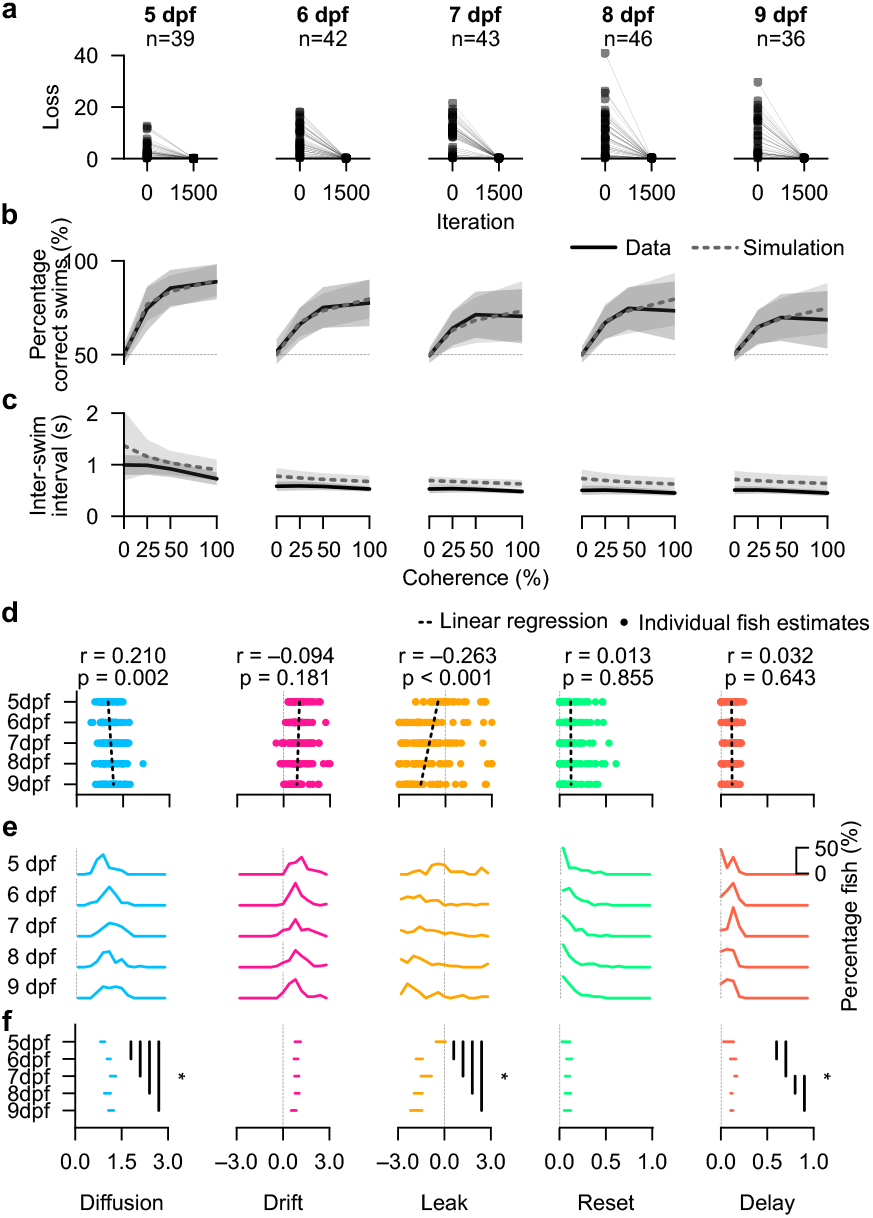
Fitting the model to experimental data reveals parameter-specific adjustments in the drift-diffusion algorithm during early development. **a**, Convergence of the loss function between the start and the end of the fitting procedure for 5 to 9 dpf. Each connected pair of dots represents one real experimental fish. Fish are not the same individuals across age groups. The number of animals per age is reported on top of each plot. **b**,**c**, Mean population accuracy (b) and inter-swim interval (c) plotted against the coherence level for experimental (solid line) and simulated data (dashed line) after fitting. The shaded areas surrounding each line represent the respective standard deviations. **d**, Individual fish parameter estimates (solid colored dots) for the different age groups together with linear regression fits (dashed black line). Regression slope and significance are displayed in the title of each panel. **e**, Distributions of the estimated individual animal parameters across age groups. **f**, Confidence interval (spanning from 5^th^ to 95^th^ quantile) of the bootstrapped median for the parameter estimations in each age group (**Methods**). For each parameter, vertical black lines indicate significantly different age groups (asterisk, p < 0.005, based on a Bonferroni-corrected threshold, corresponding to 10 pairwise comparisons – testing each age group against the other – with family-wise error rate *α* = 0.05). Sample sizes (number of fish) are given in the titles in (a).

These dynamics contrast with those observed for positive leak values. In that regime, the integrator variable decays toward zero in the absence of input, gradually forgetting previously accumulated evidence. Positive leaks formed the basis of our previous model implementations (*17, 18, 28*). The variability of the extracted parameters across fish suggests that individual larvae may employ differently tuned algorithms to integrate sensory evidence over time. Given our reliable and repeatable parameter recovery results using artificially generated data (**Fig. 2e** and **Extended Data Fig. 2b**), most of the observed variability likely reflects genuine biological differences rather than fitting imprecision. We next asked to what extent the library of optimized models could predict behavioral features not explicitly used in the fitting procedure.

To this end, we followed our previous analysis framework (*17*) and quantified swimming decision events binned across time within trials as well as on a consecutive event-to-event level (**Extended Data Fig. 4a,b**). Both experimental and simulated data showed improving accuracy over stimulus presentation time, indicating temporal integration. This observation was also true for the first swim event after stimulus onset (**Extended Data Fig. 4c**), validating that the integrator operates on temporally integrated sensory evidence. This finding also demonstrates that dynamics cannot be explained by motor memories because, for the first swim during the stimulus, previous motor events happened during the 0% coherence phase and therefore cannot provide evidence about the stimulus. In both data and model, swimming tendencies persisted for a few seconds after the stimulus turned off (**Extended Data Fig. 4a–c**). These dynamics are reminiscent of the partial resetting after each swimming decision. During periods of 0% coherence, our optimized models reproduced the tendency of animals to consecutively swim in the same direction (**Extended Data Fig. 4d)**, as previously described (*48*).

In summary, the model-based extraction of latent parameter sets from individual animal behavior provides a window into the cognitive algorithms and potential circuit mechanisms across individual animals. The close resemblance of model behavior and experimental data adds further confidence that the drift-diffusion framework can faithfully match sensorimotor decision-making in larval zebrafish and generalizes across conditions. We therefore next explored the applicability of our approach to other experimental configurations.

### Early development shapes integration dynamics

While animals develop and grow older, they face new environmental challenges, adjust their navigational goals, and thus need to integrate and interpret cues differently. Our approach of extracting drift-diffusion parameters from individual animal behavior allows us to reliably explore changes in the cognitive computations underlying such processes. For example, with increased cognitive abilities, it may be possible that animals become better at extracting signals from noise or that they display improved abilities to keep and enhance persistent internal neural representations of sensory cues, reminiscent of longer working memories. To test these ideas, we performed experiments on zebrafish larvae at ages ranging from 5 to 9 dpf. During this early period of their life, neural circuitry rapidly matures, largely extending behavioral repertoires (*49*–*51*). We thus also expect differences in their behavior to visual motion stimuli. We employed the same random-dot-motion paradigm and tracked animals in the same behavioral setups as before (**Fig. 1a,b**). Our validated fitting approach (**Fig. 2**) then enabled us to map drift-diffusion model parameters to experimental data of individual animals. Note that, due to regulatory animal permit constraints, animals need to be sacrificed after experimental sessions, and thus each age group contains different animals from the same batch.

For each age group and individual animal, our fitting algorithm reliably minimized the loss function toward near-zero values (**Fig. 4a** and **Extended Data Fig. 3h**). The loss function compares inter-swim interval distributions for correct and incorrect swims across coherence levels between model and experiment (**Fig. 3a**). Thus, the good fitting quality corroborates that a drift-diffusion model can explain zebrafish behavior for animals older than 5 dpf as well. In agreement with these results, we found the percentage of correct swims (**Fig. 4b**) and the inter-swim interval as a function of coherence levels (**Fig. 4c**) to match well between experiment and model. These results show that when animals become older, on average, they swim more frequently while making fewer correct swimming decisions in the direction of motion. What could be the algorithmic difference that gives rise to these transitions?

To answer this question, we explored the extracted parameter distributions from the fitted models across the different age groups. As a first quantification, we performed linear regression analysis for each of the extracted parameters across individuals and age groups (**Fig. 4d**). We also implemented a bootstrapping-based strategy, enabling more fine-grained statistical comparisons between each age group pair (**Fig. 4e,f**). Both analyses methods provided similar insights: Some parameters changed during development, while others remained largely stable. These findings allow us to formulate hypotheses about potential mechanistic implementations of flexibility in the cognitive algorithm:

We found the diffusion parameter to slightly increase with age, reminiscent of an increased noisiness of the decision variable while animals mature. This result may indicate that animals also start to sample other cues from the environment, in addition to motion, that they become more flexible and unpredictable in their behavioral regulation, or that their nervous system activity becomes more dynamic. The drift parameter remained stable, which suggests that the ability to extract momentary motion evidence from noisy sensory scenes does not change over early development. We observed the most remarkable parameter shift for the integrator leak. While the population distribution was centered around zero at 5 dpf, it became significantly negative starting from 6 dpf. This means that the self-reinforcing priming property of the integrator variable *x* stabilizes and becomes more prevalent across the population as animals become older. This finding may reflect increasing recurrent interactions within neural circuits supporting evidence accumulation as the nervous system matures during early development. We did not find a difference for the reset factor, making it possible that the overall mechanisms of swim event-triggered resetting of the integrator value remain unchanged. For the delay parameter, our regression-based quantification did not find an age-dependent progression. Animals at 7 dpf however had a slightly larger delay value than all other groups. While this effect was weak, it may hint at specific behavioral traits to be expressed only at short periods during development and could be linked to the onset of, for example, feeding behavior. In addition, the delay parameter seemed to be less distributed in older animals, reminiscent of an improved ability to swim more frequently with more precise motor control. These results are likely not due to batch effects, as animals of different age groups stem from the same parental crosses with different raising times (**Methods**).

Thus, our behavior-based fitting approach indicates that the drift-diffusion model can robustly capture larval zebrafish behavior across a wide range of developmental stages. The extracted parameter value distributions reveal clear adjustments of the cognitive algorithms, which may hint toward distinct modifications of their neural circuit implementations. In addition to developmental age, the genetic background of animals is likely another main driver of sensorimotor decision-making variability, an important aspect that we sought to explore next.

### Larvae with disease-related mutations have altered persistent integrator dynamics

Neurons and circuits are regulated by a complex cellular machinery, which ultimately shapes behavior. This can be clearly evidenced in several neurological disorders where mutations in one or more genes lead to devastating cascades of outcomes. However, linking specific genes and their functions to a disease phenotype has always remained a big challenge in biology.

To address this problem, we quantified the modulation of the drift-diffusion algorithm in zebrafish lines carrying knockout mutations in genes related to human autism and epilepsy (*42*), *scn1lab* and *disc1* (*18*). *Scn1lab* (*sodium voltage-gated channel alpha subunit 1a,b*) encodes a sodium channel and can be linked to neuronal excitability, while *disc1* (*disrupted-in-schizophrenia 1*) encodes a scaffolding protein that is needed for precise neurodevelopment. We previously explored these genes in the context of zebrafish social behavior and quantified responses to random-dot-motion cues (*18*). Due to the lack of reliable model fitting methods, we could previously not faithfully and automatically tune the parameters of another variant of the drift-diffusion model to the behavior (*18*). With our new validated framework (**Fig. 2**), we have now overcome this problem.

We first analyzed *scn1lab* heterozygous mutant animals *(scn1lab*_*allele1*_^+*/*–^) at 7 dpf and their homozygous control siblings (*scn1lab*_*allele1*_^+*/*+^). Larvae were genotyped after the experiment, an experimental design that removes inter-clutch variability and allows for a blinded study (see method details in ref. (*18*)). We worked on animals with heterozygous mutations because homozygous mutations are lethal early in development. For both genotypes, after model fitting, we obtained loss function values near zero (**Fig. 5a** and **Extended Data Fig. 3h**). This finding shows that the drift-diffusion modeling framework can also capture inter-swim interval distributions across coherence levels for genetically modified animals, adding further confidence to our modeling approach. In agreement with the close quantitative agreement, we observed matching between model and experimental data for the percentage of correct swims (**Fig. 5b**) and the inter-swim interval (**Fig. 5c**). We found small differences in the percentage of correct swims between the fitted model and experiment for intermediate coherence levels (**Fig. 5b**) in both groups. These differences may be explainable by small variations in the design of the random-dot-motion stimulus in our previous work compared to this study (**Methods**). We next analyzed the extracted parameter distributions after fitting. Values for the control animals (*scn1lab*_*allele1*_^+*/*+^) were qualitatively in agreement with our results for 7 dpf animals (compare **Fig. 5d** and **Fig. 4e,f**). Residual differences across our new and previously acquired datasets may originate from slightly different rearing conditions in two animal facilities and different genetic background (*scn1lab*_*allele1*_^+*/*+^ versus wild type).

**Fig. 5.**
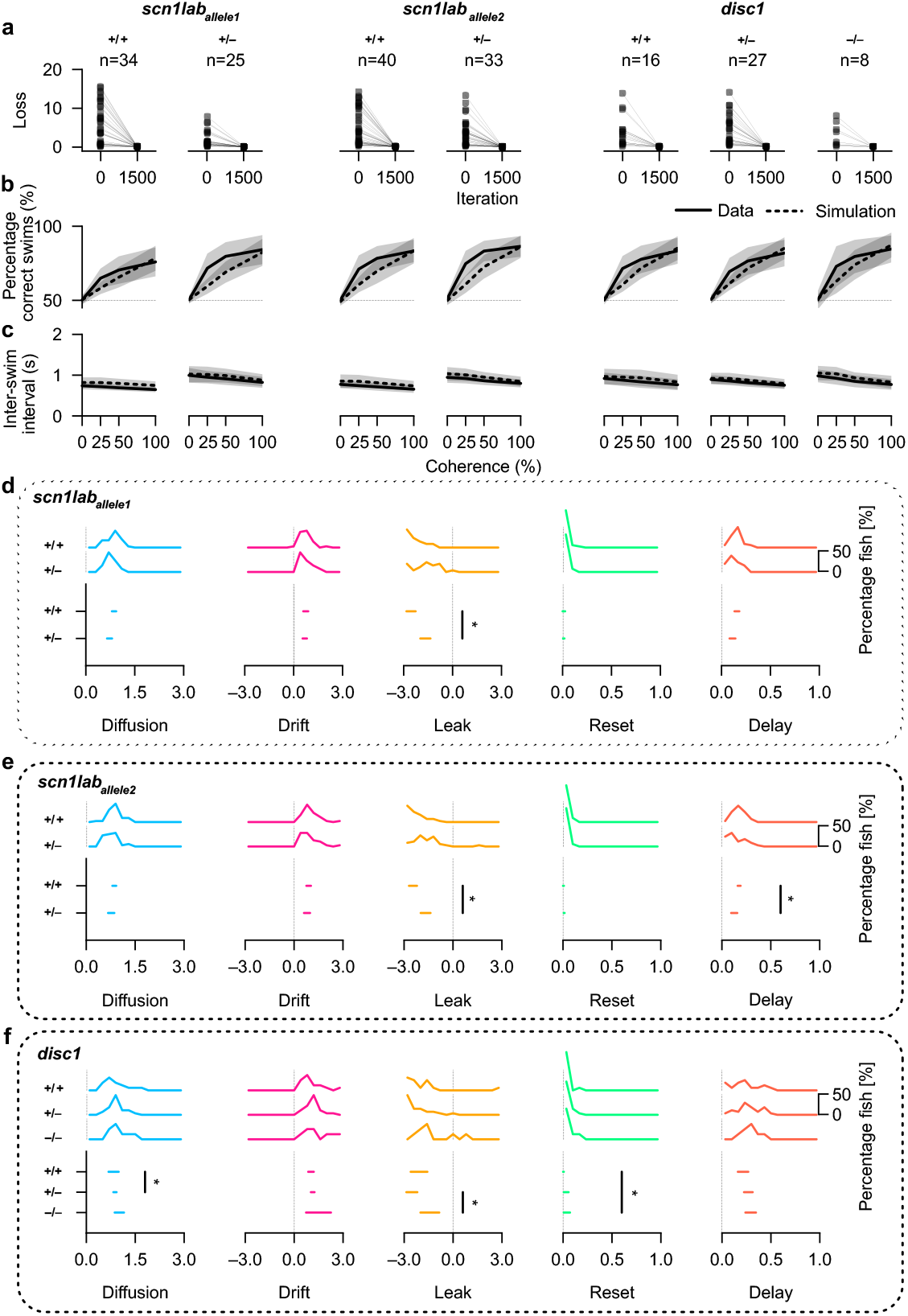
Knockout of the *scn1lab* and *disc1* genes in 7 dpf larvae reduces the self-reinforcing priming dynamics of the integrator variable. **a**, Convergence of the loss function between the start and the end of the fitting procedure for each genotype. Each connected pair of dots represents one fish. The sample sizes are presented on top of each plot for each group. We test two mutant alleles for the *scn1lab* gene (allele1 and allele2) and one for the *disc1* gene. Homozygous mutations in the *scn1lab* gene are lethal early in development for both alleles. We therefore can only test heterozygous mutants (*scn1lab*^+*/*–^) against homozygous controls (*scn1lab*^+*/*+^). As homozygous mutations in the *disc1* gene are viable, we can test heterozygous and homozygous mutants (*disc1*^+*/*–^, *disc1*^−*/*–^) against respective controls (*disc1*^+*/*+^). **b**,**c**, Percentage of correct swims (b) and inter-swim interval (c) as a function of coherence level for experimental (solid line) and simulated data (dashed line) for each genotype. The shaded areas surrounding each line represent the respective standard deviations across fish and optimized models. **d–f**, Distribution and confidence intervals (spanning from 5^th^ to 95^th^ quantile) of the estimated drift-diffusion model parameters across fish in each genotype. Confidence interval of the bootstrapped median for the estimated parameters (**Methods**). The asterisk (*) indicates a significant difference in the estimated parameter between the two genotype groups for the leak parameter (p < 0.005, bootstrapping test, based on the same Bonferroni-corrected threshold as in **Fig. 4f**. All larvae are 7 dpf old. Animals were tested blind and genotyped after the experiment (**Methods**).

Comparing mutant (*scn1lab*_*allele1*_^+*/*–^) and control (*scn1lab*_*allele1*_^+*/*+^) larvae, we found all parameters— except for the leak value—to be preserved. It thus seems that the mutation selectively acts on specific computational elements of the sensorimotor integration process. The leak parameter λ was significantly less negative in *scn1lab*_*allele1*_^+*/*–^ fish, generating weaker self-reinforcing priming dynamics of the integrator variable and potentially shorter persistent internal states. Sodium channels are key regulators of neuronal excitability, and their defects have been shown to greatly impair function and memory-like properties in neurons (*52*). Hence, our finding raises the possibility that the impaired integration dynamics in *scn1lab*_*allele1*_^+*/*–^ mutant larvae may be an emergent property of a less balanced and less recurrent neural network in the larval zebrafish hindbrain (*17, 53, 54*). The behavioral observables of these internal changes are represented by larger inter-swim intervals, with minor reductions in the percentage of correct swims (**Fig. 5b,c**), which agrees with the idea of negative-leak-dependent persistent internal states and accelerating decision threshold crossings.

To further validate our observations and to check for potential zebrafish background-specific features, we repeated our analysis in a different zebrafish line carrying another mutation in the same gene (**Fig. 5a–c**, middle, *scn1lab*_*allele2*_). Remarkably, the behavior and cognitive parameters extracted for this second allele and the differential phenotypes between mutant and control were almost identical to the results obtained in the *scn1lab*_*allele1*_ fish line. Also for *scn1lab*_*allele2*_, we found the leak parameter λ to be significantly less negative in the mutant animals compared to control conditions (**Fig. 5e**). The delay parameter turned out to be slightly lower in the heterozygous mutant group compared to the control, qualitatively matching the trend visible in the other tested *scn1lab*_*allele1*_ line (**Fig. 5d**). This finding raised the possibility that neuronal excitability through specialized sodium channels can also regulate other aspects of the mechanistic implementation of the drift-diffusion model. This complementary dataset thus adds further and independent confirmation of the potential role of the *scn1lab* gene in the regulation of neuronal excitability within decision-making circuits.

We further quantified behavior and model parameter fitting results in *disc1* mutants (*disc1*^−*/*–^) and their respective sibling heterozygous and homozygous controls (*disc1*^+*/*–^ and *disc1*^+*/*+^, respectively). Behavior and fitting quality were similar to the results obtained for *scn1lab*_*allele1*_ *scn1lab*_*allele2*_ animals (**Fig. 5a–c**, right). Across the three genotypes, only a few extracted model parameters changed significantly (**Fig. 5f**). In larvae carrying heterozygous mutations (*disc1*^+*/*–^), we observed a slight increase in the diffusion value compared to homozygous controls. In homozygous *disc1*^−*/*–^ mutants, the reset parameter was slightly more positive relative to the control. The integrator leak values displayed the strongest effect and were less negative in homozygous *disc1*^−*/*–^ mutant animals when compared to heterozygous controls. This result suggests a possible defect in the developmental arrangement of neural circuitry underlying persistent internal neural states and robust evidence integration in *disc1*^−*/*–^ mutant animals.

In summary, the application of our drift-diffusion model fitting pipeline to animals with specific disease-related mutations shows that we can generate testable experimental hypotheses for precise phenotypic analyses. Disrupting genes related to either neuronal excitability (*scn1lab*) or molecular structural pathways for development (*disc1*) leads to less negative leak values and thus less robust self-reinforcement and persistence of the integrator during the sensorimotor decision-making process.

## DISCUSSION

We show that our approach enables high-throughput, individual-level inference of latent decision variables in a vertebrate model, generating biologically interpretable insights into developmental processes and disease-associated genetic perturbations.

The drift-diffusion model provides a popular framework for the study of decision-making across species (*3, 7, 10, 45*). Despite its simplicity, the model accurately predicts response accuracy and delay across a wide range of experimental conditions. Here, we have employed the drift-diffusion model to systematically explore how larval zebrafish integrate and transform noisy whole-field random-dot-motion cues into behavioral actions. Using the optomotor behavior of larvae, we find that the inter-swim interval distributions—a proxy of response delay—shift toward shorter values for larger coherence levels. With increasing coherence, fish are also more likely to follow directional motion cues—a proxy of response accuracy. These observations are in agreement with previous experiments in larval zebrafish (*17, 28, 38*), corroborating that the optomotor response provides a tractable paradigm for studying motion-evidence integration during sensorimotor decision-making. We then fitted the drift-diffusion model to our experimental data, enabling biologically interpretable inferences about latent computational mechanisms in the brain.

A general challenge with model fitting is that extracted model parameters may depend on each other because of inadequate model design or the lack of appropriate optimization methods. Such parameter interdependencies and resulting identifiability problems are well known in cognitive modeling (*16, 25, 55*). In our previous work, we used manual parameter tuning (*17*) or complex multi-objective strategies (*18*). These approaches could not automatically extract parameters from behavioral data, limiting the systematic inference of cognitive mechanisms from experimental observations. Here, we developed a Bayesian-optimization-based pipeline to tune the parameters of drift-diffusion models to high-throughput experimental data from individual animals, enabling robust parameter recovery across repeated runs. We carefully validated our method on model-generated datasets to confirm that we can reliably and repeatedly extract parameter values from individual animal behavioral data at experimentally feasible time scales. These fitted models then enabled us to extract latent cognitive variables and to formulate testable hypotheses about their neural mechanistic implementations. Using this approach, we observed enhancements of the self-reinforcing dynamics of the integrator variable as animals mature. This feature was reduced in mutant larvae with deficits in neuronal excitability and structural development.

An important question is to what extent the drift-diffusion model class provides a good approximation of the computational processes in the brain of larval zebrafish during sensorimotor decision-making. Our loss function quantifies how correct and incorrect response delay distributions match between experiment and model. Loss values converge toward near-zero levels during the optimization across a wide range of experimental conditions, supporting that this model class captures key behavioral statistics in our task. The anterior hindbrain of the larval zebrafish has been shown to integrate random-dot-motion stimuli over time (*17, 38*), making it a candidate brain region involved in the integration of sensory evidence. Together, these results establish larval zebrafish as a uniquely accessible vertebrate system for mechanistic dissections of how neural circuits may implement evidence integration algorithms within a drift-diffusion framework.

Our study currently relies on binarized leftward and rightward swimming events, even though turning behavior follows a distribution of angular changes. This simplification facilitated interpretation and model design, but future work could extend our approach to incorporate more graded or confidence-like readouts of decision-making (*56*). The decision bound variable was mathematically redundant, scaling with the other parameters that drive the integrator variable. As such, we treated the decision bound as a fixed parameter and did not include it in the fitting procedure. However, animals may dynamically regulate decision bounds depending on task demands, such as time pressure or stimulus context (*57*–*60*). Allowing dynamic bounds would provide a more complete description of the decision-making algorithm in our random-dot-motion paradigm. While our current parameter recovery performance is robust and yields biologically interpretable features, expanding the model to include additional free parameters will likely necessitate further improvements to distance metrics and possibly more advanced optimization methods.

Several forms and variants of drift-diffusion models have been adopted to describe decision-making processes over the past few decades (*8*). For example, the integrator variable may follow a random walk or diffusive process, it may leak toward zero, or multiple competing integrators can operate simultaneously (*46*). In larval zebrafish, it has been proposed that animals may employ a stochastic readout of the integrator variable with different rates above or below the threshold (*17, 18*), and that they occasionally lose attention to the stimulus (*28*). These models are more complex variants of the model used in this work, which makes automatic fitting approaches substantially more challenging. Which of these model variants best approximates the computations implemented by the larval zebrafish brain remains to be determined and will require systematic mapping of the proposed computational processes onto neural representations. Here, using the simplest possible model that can be reliably and automatically fitted to the behavior of individuals is therefore a good starting point for generating experimentally testable hypotheses.

Fitting drift-diffusion models to behavior has a long tradition in neuroscience, in particular in human psychology, allowing for principled inference and interpretation of latent variables from experimental data (*15, 26, 27, 61*). In rats, fitting drift-diffusion models to auditory discrimination tasks has shown that animals can solve tasks with near-optimal performance (*2*) and that latent accumulation variables have identifiable neural correlates even across trials (*62, 63*). The rat parietal and frontal cortices have been shown to have differential roles (*64, 65*). Drift-diffusion models have also been successfully employed to describe foraging behavior in rodents, allowing discrimination of distinct search strategies (*44*). In *Drosophila*, drift-diffusion processes have also been attributed to categorical choice in odor discrimination tasks, identifying specific mushroom body compartments to implement these processes (*5, 45*). Moreover, in flies, the landing response to repeated stimuli has been modeled through an integration process (*4*).

All these studies assume that the drift-diffusion model is a sufficiently good approximation of the computational processes in the brain of animals, which may not always be the case. Other work has proposed that drift-like integration dynamics may not necessarily arise within individual trials, but may instead be an artifact of trial-by-trial averaging (*24*). These findings caution against equating good model fits with direct neural implementation and suggest that one must carefully evaluate hypotheses about potential cognitive mechanisms through direct observational and causal experimentation.

Larval zebrafish offer the unique advantage of relatively easy whole-brain functional imaging to map model computations to real neuronal implementations (*17, 37, 39, 66, 67*). For example, we found that the leak parameter gets progressively more negative while animals mature. In our model, a negative leak implies that accumulated evidence is self-reinforcing, maintaining elevated states even without ongoing sensory input (*68, 69*). A negative leak has also been observed in rat decision-making studies and might support faster or more robust decisions through recurrent network states (*1, 2, 64*). In the developing larval zebrafish, we hypothesize that such dynamics may expand the behavioral repertoire in more challenging sensory environments, in particular when animals grow older and start to navigate and forage for food. It would be interesting in future experiments to perform functional imaging of known structures with integration dynamics and persistent activity, such as the anterior hindbrain (*17*), and compare dynamics across different developmental stages.

Furthermore, our explorations also predicted that the dynamics of the integrator variable should change in fish carrying mutations in the genes *scn1lab* and *disc1*, which affect neuronal excitability and neurodevelopmental circuit structure, respectively (*42*). Neuronal excitability, neurotransmitter balance, and recurrent network configurations are tightly linked to implementing slow temporal dynamics, neural state persistence, and working memories in the brain (*68*–*70*). Therefore, disrupting such arrangements in mutant larvae may contribute to the observed increase in integrator leakiness. Increases in leakiness reduces the rate of swimming with only modest effects on the correctness of sensorimotor decision-making. In the brains of these animals, one would expect less self-reinforcing or faster decaying response dynamics of integrator cells during random-dot-motion. Such results may then point toward potential mechanisms of neuronal dysfunction in neuropsychiatric conditions, with which these genes have been associated in humans (*71, 72*). Our work of extracting latent cognitive model variables could thus also provide a novel and powerful approach for more precise high-throughput phenotyping in large-scale mutant or drug screens.

Our behavior-based latent parameter estimation approach provides a window into the hidden dynamics governing sensory evidence accumulation in a vertebrate model system amenable to high-throughput behavioral phenotyping, genetic screens, and brain-wide neural imaging. Our framework only requires observable event times and correctness labels and is thus readily extendable to decision-making paradigms across model systems, including insects, rodents, primates, and humans, where drift-diffusion models have been widely used to describe behavior.

Future functional brain imaging or causal manipulation experiments may test—and potentially falsify—the predictions of our behavior-constrained drift-diffusion models. In such cases, our optimization approach can be updated and extended, and parameter estimation procedures can be repeated accordingly. Through an iterative process of neural mechanistic analysis, model refinement, and circuit interrogation, this approach can converge on increasingly precise descriptions of the sensorimotor decision-making process and facilitate the dissection of underlying neural circuits. Establishing such a model–experiment–model loop is a central strategy in systems neuroscience to link persistent network dynamics to behavioral observables, and our paradigm thus provides a scalable framework applicable across species and experimental systems.

## MATERIAL AND METHODS

### Animals

All experimental data were collected from zebrafish larvae aged 5 to 9 days post-fertilization (dpf). Experiments were approved by the Regierungspräsidium Freiburg under permit number G21/153. Larvae originated from group-crossed wild-type AB adults. Eggs were collected in the morning, within a few hours from fertilization, and immediately transferred to large Petri dishes (14.5 cm diameter) containing standard 1x E3 fish water (5 mM NaCl, 0.18 mM KCl, 0.33 mM CaCl_2_, 0.4 mM MgCl_2_) (*73*) supplemented with methylene blue. Each dish contained a maximum of 50 eggs and was placed in an incubator (Memmert IPP260eco) running at a constant temperature of 28°C and a 14/10-hour light/dark cycle.

Before reaching 3 dpf, larvae were transferred to fresh E3 medium without methylene blue. Water was replaced daily in the morning. For the experiments on wild-type fish ranging from 5 to 9 dpf (**Figs. 1, 3**, and **4**), larvae were fed powdered food (Sparos ZEBRAFEED < 100 μm diameter) twice a day, starting at 5 dpf, once between 9 and 10 AM and again between 3 and 4 PM. For the experiments in **Fig. 4**, 16 larvae were randomly selected from the Petri dishes each afternoon and transferred to the experimental arenas. To avoid potential batch effects, we used offspring from the same parental crosses, raised over different numbers of days. We performed these experiments over multiple crosses and weeks, merging offspring from different parents. Larvae were allowed to acclimatize for 30 minutes in the arena before the experiment. Animals were sacrificed immediately after the experiment through rapid chilling. Animals carrying disease-related mutations (**Fig. 5**) were raised similarly but in another zebrafish facility (18). See details below.

### Behavioral tracking and swimming event analysis

Behavioral data are acquired in parallel by multiple custom-built high-throughput behavioral tracking setups (**Fig. 1a**). We used the setup design as previously reported (*74*). Briefly, each setup supports the simultaneous acquisition of eight fish at a time, each freely swimming in a separate custom-made arena dish. Every dish is associated with a dedicated camera, centered on top of the arena. Stimuli are projected from below via AAXA P300 Pico projectors. The bottom of the dishes was coated inside the water with diffusive paper to distribute projector light evenly. Dishes had a radius of 6 cm and were surrounded by a black acrylic rim.

To prevent the stimulus light from interfering with tracking, infrared-sensitive cameras (Basler acA2040-90um-NIR) were equipped with a long-pass infrared filter (Linghuang Zomei IR 850 nm 52 mm). Infrared light at 940 nm was provided through a custom-built matrix made of LED strips at the bottom of the behavioral setup. Thus, only the infrared shadow of the fish is visible to each camera. Cameras acquired frames at 90 Hz, with incoming data being processed in real-time. This configuration enabled closed-loop experiments, such that stimuli could be directly linked to the behavior of the animal.

Animal posture was calculated using a custom-written posture analysis from the largest identified contour after background subtraction. Heading orientation was then determined for each frame by computing the principal component vector of the head. The time trace of orientation was then analyzed through a rolling window variance with a window size of 50 ms. Swim event starts were defined as when the variance exceeded 1 deg^2^ for at least 20 ms. Swim events ended when variance fell below 0.5 deg^2^ for at least 50 ms. The tracking system then stored the timing of swims and by how much animals turned per swim. These data were used to calculate all metrics presented. The inter-swim interval describes the time span from the end of the swimming event to the beginning of the next swimming event (**Fig. 1c**). The distribution of heading angle changes during swims (**Fig. 1f**) then allowed us to assign correct or incorrect labels, depending on whether animals turned in the direction of coherent motion or went against it, respectively. Forward swims between ±3 deg were discarded (around 19% of all swims in the 5 dpf dataset). The binarization of swim events substantially simplified all subsequent experimental analyses and computational modeling.

To remove potential tracking errors, the following quality control metrics were applied: 1) inter-swim intervals needed to be below 30 s. Fish usually swim once a second on average; much longer intervals indicate unhealthy animals or extended periods of quiescence. 2) The distance divided by the duration of the swim (average swim speed) needed to be below 6 cm/s. Longer values are considered tracking errors, where the algorithm finds other structures in the arena. 3) Fish contour areas had to be smaller than 2000 pixels. Larger values are likely accidentally tracking issues of the arena rim. 4) The turn angle of each swim should not exceed 150 deg. Such large values are unrealistic and indicate events where the tracking system accidentally swaps head and tail. 5) If more than 5% of swims were considered tracking errors by any of these criteria, the entire trial was dropped. Such periods likely indicate animals to be too close to the wall for longer periods or that the system has started to track dust particles in the water. 6) We discarded animals when swim rates were, on average, below 0.375 swims/s (around 19% of larvae in the 5 dpf dataset and around 4% in all other datasets). Such low average rates point to unhealthy or underdeveloped animals.

### Visual stimuli

The stimulus used in our experiments was a random-dot kinematogram (**Supplementary Movies 1– 3**). Dot movement direction was updated in real-time so that dots continuously moved rightward (– 90°) or leftward (+90°) relative to the body orientation of the freely swimming fish (**Fig. 1b**). This configuration ensured that stimuli always remained perpendicular to the animal and in the same direction within a trial.

The implementation of the stimulus was slightly different from what we used previously (*17, 28*), better matching classical variants (*6*). Specifically, we now define a frame-to-frame update rate of 30 Hz at which dots get redrawn. During each trial, at every frame update, all dots are randomly divided into two groups: the coherently moving group and the randomly moving group. The coherence level of the trial determines the percentage of dots assigned to the first group, while the others fall into the second group. Dots belonging to the first group are then placed spatially offset by 0.6 mm, while those in the second group are redrawn at a random location in the arena. Thus, at a 100% coherence level, all dots move coherently in one direction without flickering. At intermediate coherence levels, dots most often only move for one or two frames before being randomly repositioned. For 0% coherence stimuli, all dots are randomly redrawn at 30 Hz. Dots were small (∼2 mm diameter) and light gray on a black background. We used 1,200 dots per arena. The overall brightness of the scene was 61±7 lux (LX1330B light meter, Hongkong Thousandshores Ltd., Central Hong Kong). All stimuli were rendered online using custom-written software implemented in Python 3.12 and Panda3D 1.10 with OpenGL Shading Language (GLSL) vertex shaders running on AMD Radeon RX 580 GPUs.

Each experimental session lasted for two hours and consisted of repeated trials with a fixed temporal structure: 10 seconds of rest (0% coherence), followed by 30 seconds of stimulus (0%, 25%, 50%, or 100% coherence), and another 10 seconds of rest (0% coherence) (**Fig. 1e**). Both the movement direction and the coherence level were randomly selected for each trial. During the presentation of the 30-second-long stimulus, animals swam multiple times (**Fig. 1e**). All these events were considered in our analysis (see above).

### Experiments with larvae carrying disease-related mutations

For the experiments with larvae carrying disease-related mutations (**Fig. 5**), we used published data from our previous work (*18*). This dataset includes inter-swim intervals and correct and incorrect decision labels for control homozygous wild-type *scn1lab*^+*/*+^ and heterozygous mutant *scn1lab*^+*/*–^ fish. We used two alleles, one fish line that had previously been generated at the Novartis Institutes for BioMedical Research (NIBR) in the lab of Mark C. Fishman (*42*) and one fish line from the Zebrafish International Resource Center (ZIRC). We named these alleles *scn1lab*_*allele1*_ (for the line from NIBR) and *scn1lab*_*allele2*_ (for the line from ZIRC), respectively. Homozygous mutations of the *scn1lab* gene are lethal early in development. The *scn1lab* gene encodes a sodium channel (Nav1.1) and is associated with autism and epilepsy (*41, 75*). Our dataset also includes homozygous wild-type *disc1*^+*/*+^, heterozygous mutant *disc1*^+*/*–^, and homozygous mutant *disc1*^−*/*–^, which had also been generated previously at NIBR (*42*). The gene *disc1* stands for *disrupted-in-schizophrenia 1* and is associated with schizophrenia, bipolar disease, and major depressive disorder (*72*).

All animals in this dataset are 7 dpf old. Experiments were performed blindly in a sibling-controlled manner. Offspring of heterozygous mutant parents were genotyped after the experiments. The experimental setup was near-identical to that used in the present study. Each trial consisted of 10 s of rest, followed by 10 s of stimulus (0%, 25%, 50%, or 100% coherence) and 5 s of post-stimulus rest. During rest, animals saw 0% coherence. During the stimulus, dots either moved rightward or leftward relative to the body orientation of the fish. Here, random-dot-motion kinematograms were implemented by assigning dots to the coherently moving group only at the start of a trial. Every dot, independently of the group to which it belonged, had a short lifetime (200 ms mean) and stochastically disappeared and immediately reappeared at a random location.

### Drift-diffusion model

We implemented the drift-diffusion model using an Ornstein-Uhlenbeck process:

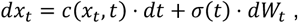

Where

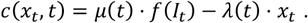

Here, *x*_*t*_ defines the decision variable, and *σ*(*t*), *µ*(*t*), and λ(*t*) represent the diffusion, drift, and leak parameters, respectively. Note that despite the general formulation allowing for these parameters to be time-variant, we will treat them as constant values within a model. *dW*_*t*_ is a Wiener process, a white noise contribution implemented using NumPy’s Gaussian random number generator. The random variable was scaled by 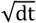 to enable comparable simulation results when using different *dt. f*(*I*_*t*_) is a function that transforms the stimulus strength *I*_*t*_ into the input to the integrator. *I*_*t*_ is a scalar value in the range 0 ≤ *I*_*t*_ ≤ 1. Following Weber’s Law of sensory processing (*67, 76*), the mapping was implemented as a sublinear transformation:

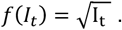

We fixed the exponent of this power law to 0.5 (= square root) to reduce the degrees of freedom of our model. When leaving the exponent parameter open during the fitting process, Bayesian optimization generated different results with repeated optimization runs. We used absorbing boundaries at ±*B*, meaning that a decision event is defined when the decision variable *x*_*t*_ hits one of the two boundaries (*x*_*t*_ V≥ *B*). The decision variable is then partially reset:

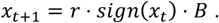

The reset factor *r* was constrained between 0 ≤ *r* ≤ 1. After each decision, we stopped the integration process for a short delay period *δ* by clamping *dx*_*t*_ to 0 for *t* < *δ*.

We solve these equations numerically by explicit forward Euler simulations for the trajectory *x*_*t*_ using Python Numba. An appropriate choice of the *dt* value is important for numerical stability and speed of the computation. A small *dt* largely extends stimulation times. We found no benefit for *dt* < 0.01*s* but considerable instabilities for *dt* ≥ 0.1*s* (**Extended Data Fig. 1e,f**). We therefore used *dt* = 0.01*s* in all other simulations in our manuscript. For each decision event, we record the time passed since the last event to obtain the inter-swim interval. We labeled events as correct or incorrect depending on whether *x*_*t*_ reaches +*B* or −*B*, respectively. This structure allows us to analyze results from simulations in the same way as experimentally obtained data, largely simplifying fitting procedures.

Notably, in the formulation of our model, the boundary is a redundant parameter that linearly scales with other integrator variables, like the drift and the diffusion. Therefore, we fixed the boundary to *B* = 1. Our model thus has a total of 5 free parameters (diffusion *σ*, drift *µ*, leak λ, delay period *δ*, and reset factor *r*).

### Distance metrics and fitting procedure

To estimate latent variables of integration, we only used steady-state experimental data and simulation results 2 s after stimulus start, excluding adaptation processes right after stimulus onset. We computed histograms of inter-swim intervals for correct and incorrect decisions for each coherence level, using a bin size of 0.05 s. We chose the bin size to be 5 times larger than our simulation *dt* to ensure robust binning of model-generated data. We then normalized histograms by the total time of stimulation over the entire experiment or simulation run. Our fitting process aims to match distributions between model and experiment (**Fig. 3a**). To robustly compute error functions, ignoring outliers, we only considered inter-swim intervals in the range between 0 and 2 s. This range includes 92.5% of the swim events for 5 dpf fish. We computed distances *d* between corresponding distributions using a weighted version of the Kullback-Leibler divergence:

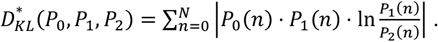

As 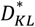 is an asymmetric metric when comparing target with model and model with target, we computed distances twice:

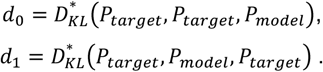

We then selected the larger value to define distances. We computed distances for each coherence level and correct and incorrect labels, resulting in eight distance values (**Fig. 2d**). To obtain a final distance value between the model and the target, we averaged all values.

We also compared results with the standard, non-weighted form of the Kullback-Leibler divergence:

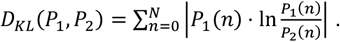

Our adaptation to the Kullback-Leibler divergence 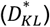 emphasizes the peak of the target distributions, more strongly weighing this part rather than flat tails. We found 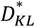 to perform generally better than *D*_KL_ (**Extended Data Fig. 1b–d**).

We adopted the Python scikit-optimize 0.8.1 implementation of Bayesian optimization (*77, 78*), using *gp_minimize* with default parameters. Specifically, the base estimator was a Matern kernel, and the initial points were generated randomly. We used *gp_hedge* as an acquisition function, which probabilistically chooses among lower confidence bound, negative expected improvement, and negative probability of improvement at each iteration. We used *lbfgs* as an acquisition optimizer. We randomly initialized the Gaussian process estimator with 100 rounds of simulation and computation of the loss, and then minimized the distance between the model and the target through 1,400 optimization steps, for a total of 1500 function calls. At each iteration, we simulated the model for 2,000 trials to obtain smooth distributions. Parameter search ranges were set based on a trial-and-error approach, such that the optimization algorithm could fit data for larvae at 5 dpf without saturating parameters at their search boundaries. We then used the selected ranges for all experimental conditions throughout the manuscript. We ran all model optimizations in parallel on the Baden-Württemberg high-performance compute cluster (bwForCluster NEMO 2) with 1 requested CPU core and a generous 64GB RAM per model. On average, it took around 20 hours for a model optimization to complete.

### Criteria for selecting target models

To generate model-based data, we chose different target parameter sets. These parameters could not simply be selected randomly, as many combinations do not lead to any threshold crossing events or do not generate biologically realistic behavior. Therefore, simulation results needed to fulfill a list of acceptability criteria:

We need to set a lower boundary to the activity at each condition, granting the model some degree of activity. Thus, the areas under the normalized histograms for correct (*A*^*correct*^) and incorrect (*A*^*incorrect*^) swims needed a minimum level of activity that scaled with coherence level *c* :

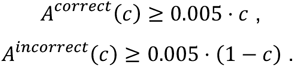

As in experimental data, we observe high activity at maximum coherence level (*c* = 1), either or both areas also had to be larger than a minimal value:

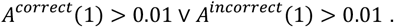

Furthermore, areas had to increase with coherence levels, and at the highest coherence level, there had to be more correct than incorrect swims:

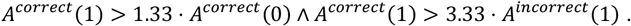

To enable further behavioral flexibility, models were also accepted if they produced more incorrect swims with increasing coherence and if there were more incorrect swims than correct swims at the highest coherence level:

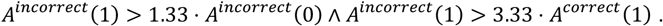

Finally, we sought to limit the value of each time bin in the histograms, thus avoiding models that generate events at an unnaturally regular frequency. We did this by only including models with realistic occurrences (*h*^*correct*^ and *h*^*incorrect*^) in any inter-swim interval bin (*τ*) and coherence level (*c*):

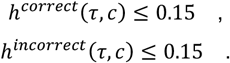

Following all these criteria, we generated a list of 100 valid target models (**Extended Data Fig. 2a**) that we could use to probe our optimization algorithm.

### Calculation of parameter estimation errors

Our optimization algorithm seeks to retrieve latent variables by fitting models to target data. For the model-generated target data (**Fig. 2e** and **Extended Data Fig. 2b**), we quantified the parameter estimation error *e* (*j*)over iterations by comparing the estimated parameter *p*(*j*) at optimization step *j* with its target value *p*^*target*^ :

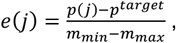

where *m*_*min*_ and *m*_*max*_ are the lower and upper search space boundaries, respectively.

For real experimental target data, we do not know the target parameter set and therefore cannot compute an error. To still be able to quantify the dynamic evolution of the optimization process, we computed the parameter estimation distance *d* (*j*) by comparing the estimated parameter *p*(*j*) at the optimization step *j* with its final estimated value *p*^*final*^ at the last optimization step:

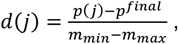

where *m*_*min*_ and *m*_*max*_ are the lower and upper search space boundaries, respectively.

### Parameter extraction quality

To probe to what degree latent parameters can be repeatedly identified from observable inter-swim interval data, we generated 100 model-generated datasets using validated target models (see model selection criteria above). We simulated each of them for a duration of 3,600 s (360,000 time steps at *dt* = 0.01*s*), matching the overall time real fish are tested in our behavioral assay. After 1,500 optimization steps, error metrics converged to near-zero (**Extended Data Fig. 2b**). For all tested target models, all estimated parameters converged to their target values (**Extended Data Fig. 2b**).

For the validation tests in **Extended Data Fig. 2c**, we simulated different experimental durations ranging from 120 s to 12,000 s and statistically compared results with non-optimized random model initializations. This analysis allowed us to define the minimal required dataset size for reliable model fitting as 1,200 s. In our experiments, we repeat 30 x 30-long trials for four coherence levels (900 s per coherence level; 3,600 s total), which should thus be sufficiently long to estimate parameters.

To probe the dependence of parameter extraction quality on the degree of noise in the time-normalized inter-swim interval distributions in synthetic data (**Extended Data Fig. 3a**), we added Gaussian noise *N*) of different variance levels *N*(0, *σ*^2^) to each bin. For experimentally realistic noise levels, parameter estimation remained robust.

### Statistics

The coefficient of variation (*CV*) across coherence levels (**Extended Data Fig. 1a**) was calculated as

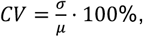

where *σ* and μ are the standard deviation and mean, respectively, across fish.

In **Figs. 4** and **5**, we used custom-implemented bootstrapping hypothesis testing to quantify significant differences across all studied conditions. To this end, we first computed the median distance between the estimated quantities in two conditions. Then, we produced two sets of datasets, pulling samples with replacement from both the original datasets 10,000 times, under the null hypothesis that there is no difference between conditions. We then computed the p-value as the percentage of cases in which the distance of the median of a pair of resampled datasets matched or exceeded the observed distance between conditions. The resulting p-value does not depend on the number of iterations, but becomes more robust for larger iteration numbers. The reported significance values were determined based on the error rate 0.05 and Bonferroni-corrected for 10 pairwise comparisons among all five tested age groups in **Fig. 4**, resulting in a threshold value of p < 0.005. While this threshold was chosen for the experiments in **Fig. 4**, we kept this relatively strict value for assigning significance also in **Fig. 5**. In **Figs. 4f** and **5d–f**, bootstrapping was also used to provide a central point of the parameter distributions in the form of 5^th^ and 95^th^ confidence intervals for the median.

We calculated statistical significance in **Extended Data Fig. 4b** by comparing the observed difference across the percentage of correct swims across consecutive decision events with 10,000 differences obtained from resampling the mixed population. We chose a significance threshold of p < 0.05.

To compute the linear regression in **Fig. 4d**, we used the standard functionality of Python SciPy (version 1.14.1) function *linregress*.

### Sensitivity analysis

To understand the impact of estimation error of each parameter on the loss function, we computed the distance between data generated from a model with randomly sampled parameter sets and those generated by the same model after perturbing each parameter independently (**Extended Data Fig. 3i**). Perturbation ranges from –50 to +50% of the search space width in each parameter. This sensitivity analysis was run on the models estimated from the wild-type 5 dpf fish (**Fig. 3f)**. As a measure of distance between datasets, we computed the loss function 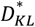 (see above) between the resulting inter-swim interval distributions. Models were not evaluated if the perturbed parameter value exceeded the boundaries of the corresponding search space, resulting in fewer data points for larger perturbations or for parameters often lying close to the edge of their search space. A Mann-Whitney U nonparametric test was used to distinguish distributions of loss values obtained for each tested perturbation level.

## Acknowledgments

We would like to thank all the members of the Bahl lab and the other Konstanz Neurobiology groups for fruitful discussions throughout this project. We thank Ulrike Bonitz and Heike Naumann for administrative support and all animal caretakers at our central fish facility for their continuous help. Furthermore, we thank Mariachiara Squillaci, Letizia Garza, Lidia Rossi, Ashrit Mangalwedhekar, Flutura Shabani, Katja Slangewal, Daniel Hummel, Margherita Zaupa, Meha Jadhav, Sophie Aimon, and Katrin Vogt for reading and commenting on the manuscript draft.

## Author contributions

Conceptualization: R.G., A.H., A.B.; Data curation: R.G., A.B.; Formal analysis: R.G.; Funding acquisition: A.B.; Investigation: R.G.; Methodology: R.G.; Project administration: A.B.; Resources: R.G., A.B.; Software: R.G.; Supervision: A.H., A.B.; Validation: R.G.; Visualization: R.G.; Writing – original draft preparation: R.G., A.B.; Writing – review & editing: R.G., A.H., A.B.

## Data and materials availability

All the source code to run analyses and simulations, as well as to generate plots, is available at the KonData repository with the persistent identifier https://doi.org/10.48606/rbfa7tt6aez1pjtj. All raw and preprocessed data originally generated for this manuscript can be found at the KonData repository with the persistent identifier https://doi.org/10.48606/kn59u9atf99nejfb. Requests for further information and resources should be directed to and will be fulfilled by Armin Bahl.

The above-mentioned persistent identifiers will only work upon final paper acceptance. For the review process, please use the following links:

Code: https://github.com/bahl-lab-konstanz/garza_et_al_2026

Data: https://cloud.uni-konstanz.de/index.php/s/ynEzXFqbp3mZeoW

## SUPPLEMENTARY MATERIAL

**Extended Data Fig. 1.**
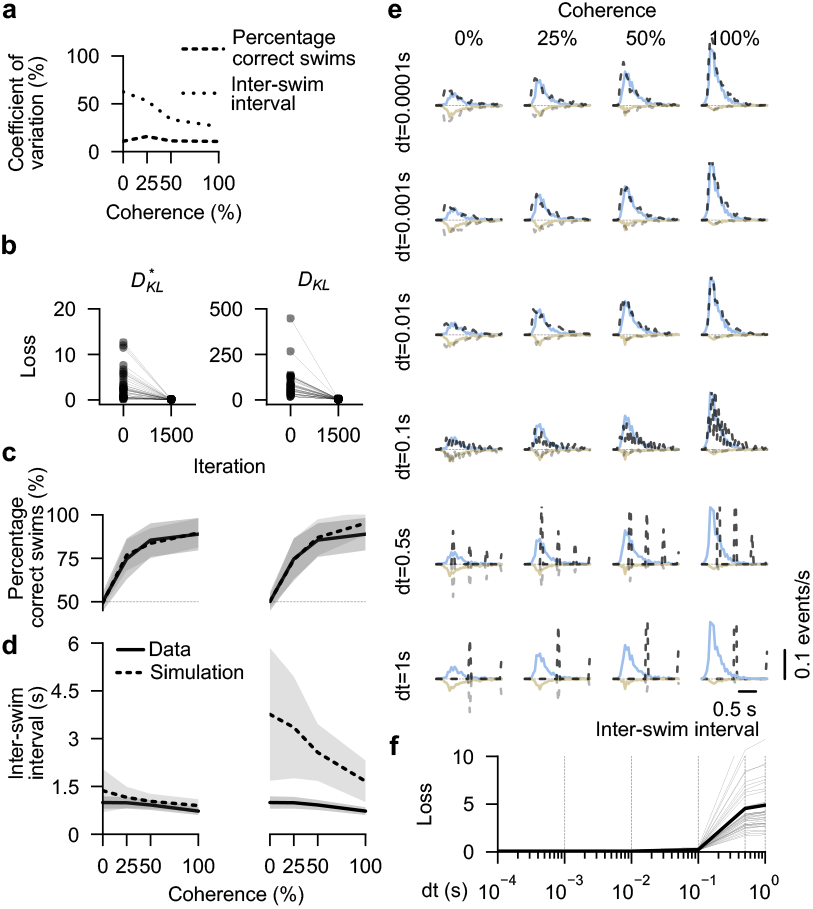
Quantification of behavioral variability. Optimization results with two different loss functions and simulation time steps. **a**, Coefficients of variation of percentage correct (long dashed) and inter-swim interval (shorter dashes) for different coherence levels of larvae from **Fig. 1g. b**, Loss function reduction before and after fit of experimental data from N=39 5 dpf real larvae, using the improved 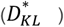 and standard (*D*_KL_) forms of the Kullback-Leibler divergence (**Methods**). Despite their difference in initial value, both loss functions converge to near-zero. **c**,**d**, Percentage of correct swims and inter-swim intervals plotted against the coherence level for the experimental (solid line) and fitted model (dashed line). The shaded areas surrounding each line represent the respective standard deviations across fish and fitted models. Optimization using 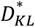 leads to better matches of the target experimental data (left side) compared to results obtained using *D*_KL_ (right side). **e**, Model fits using different simulation time resolutions ***dt*** (labeled on the left side of each row). The target dataset was always the same experimental real fish. **f**, Final loss as a function of *dt* for all N=39 5 dpf real fish. Thin gray lines indicate single fish; the thick black line is their average. The final loss stably remains near zero for ***dt*** ≤ 0.01*s*. All fish are the same individuals as in **Fig. 1g**. Related to **Figs. 1** and **2**.

**Extended Data Fig. 2.**
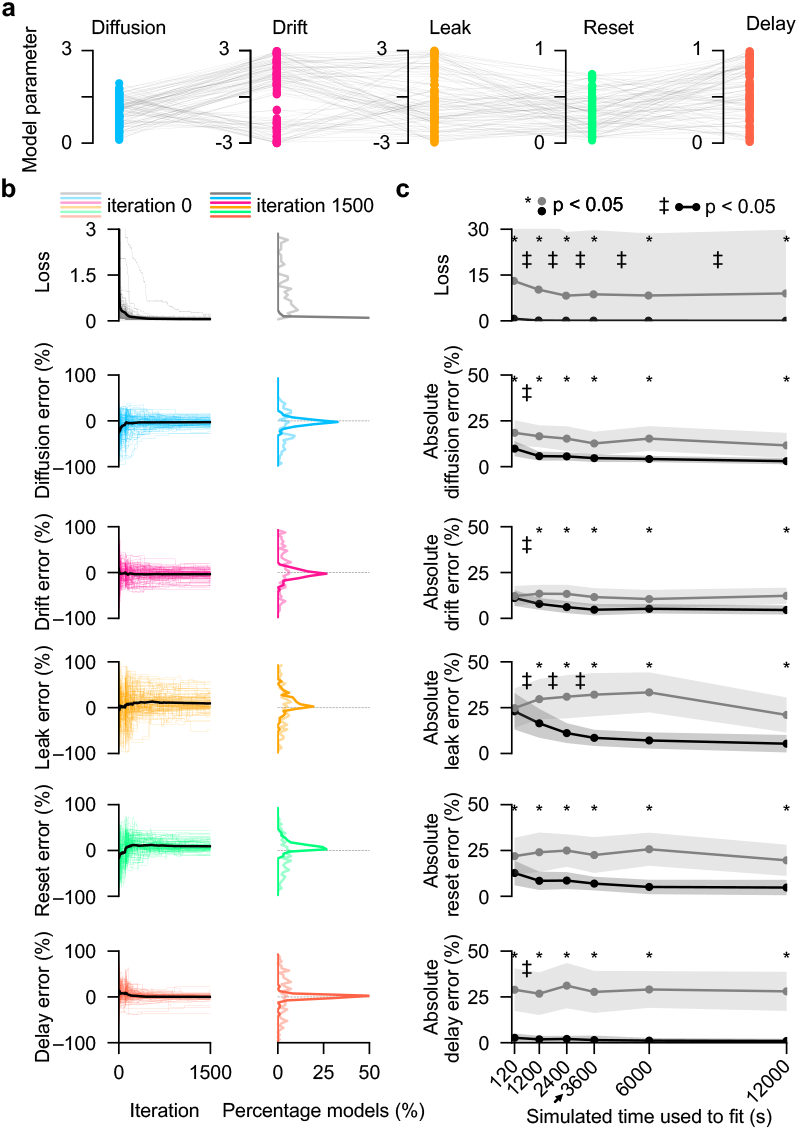
Bayesian optimization can extract latent model parameters from a broad range of datasets using experimentally feasible time scales. **a**, Parameter sets of 100 randomly generated models. These models fulfill a list of criteria to make them biologically realistic, but are not fit to experimental data (**Methods**). Gray lines connect values of the parameters belonging to the same model. **b**, Left: Evolution of loss and parameter estimation error over optimization iterations. Colored thin lines represent individual model fits. The black line shows the median across models. Right: Distribution of values at the beginning of (light colors) and after (dark colors) optimization. **c**, Loss and absolute parameter estimation errors obtained for datasets of different simulation lengths. For each length, we used the same 100 randomly generated models as in (a). The black line represents the average value across models. The black shaded area represents the corresponding standard deviation. As a control, in gray, the same results are shown for randomly initialized models before optimization. Asterisks (*) indicate significant differences (p < 0.05), comparing randomly initialized values to optimized ones. Double crosses (‡) display significant changes (p < 0.05), comparing optimized results from one simulation length to the next. Bootstrapping hypothesis testing was used in both cases. The arrow points to the size of experimental datasets. Related to **Fig. 2**.

**Extended Data Fig. 3.**
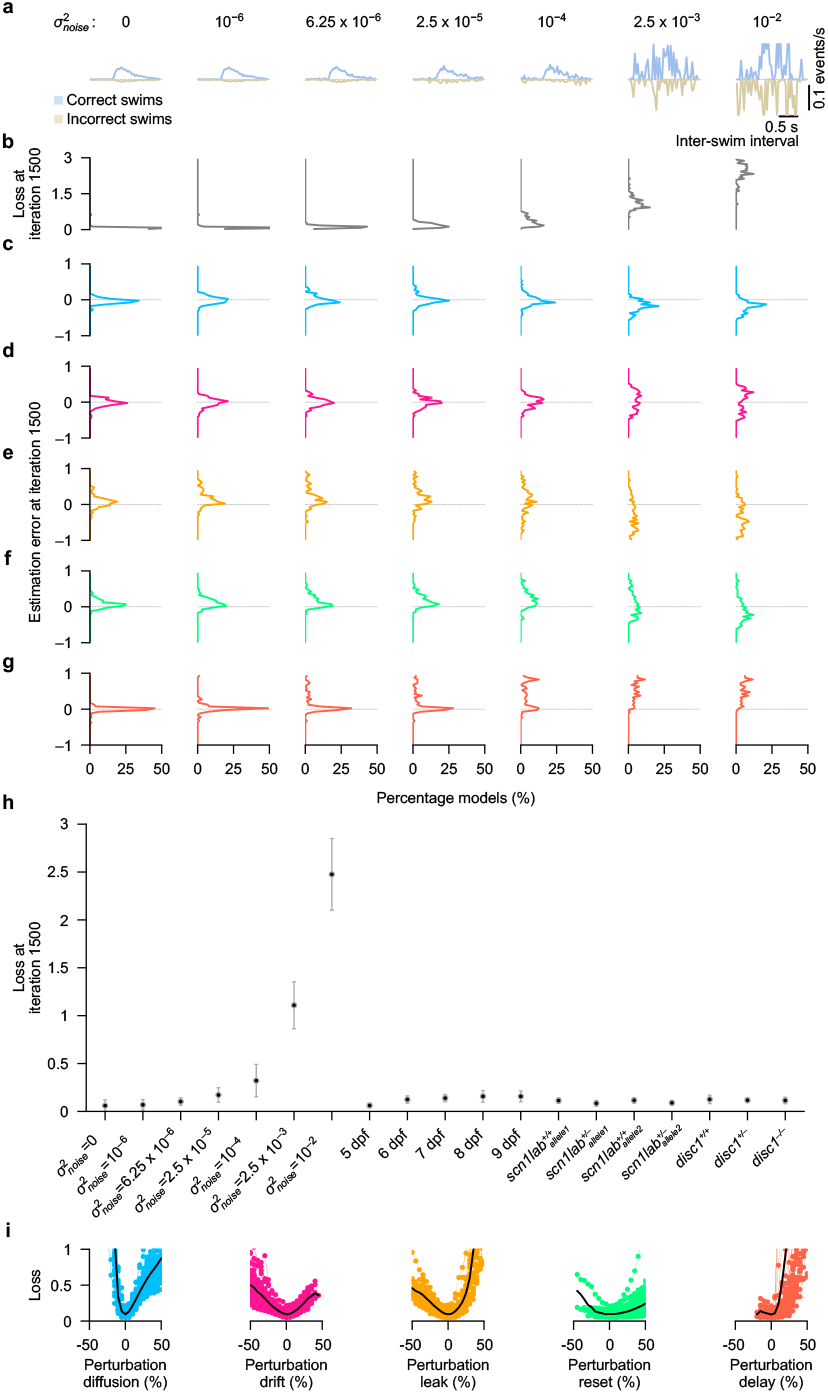
Bayesian optimization is robust to noise and generalizes across experimental conditions. **a**, Distributions of inter-swim intervals for correct swims (blue) and incorrect swims (gold) at 50% coherence from an example simulated model. We added different amounts of Gaussian noise to the distributions. 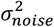 indicates the noise variance. **b–g**, Distribution of loss function value and parameter estimation errors after optimization for each noise level for 100 randomly generated models. Same models as in **Extended Data Fig. 2a. h**, Average loss function value after optimization for simulations with added Gaussian noise, quantified from (b), as well as for various experimental conditions tested in the paper. Error bars represent standard deviation. **i**, Change of loss for small parameter perturbations after fitting models to experimental data from N=39 5 dpf real larvae (same individuals as in **Fig. 1g**). Perturbations are measured in percentage of the width of each parameter’s search space. Thin colored lines link individual loss estimations (colored circles) of specific perturbation percentages for each fish and parameter. Related to **Figs. 2–5**.

**Extended Data Fig. 4.**
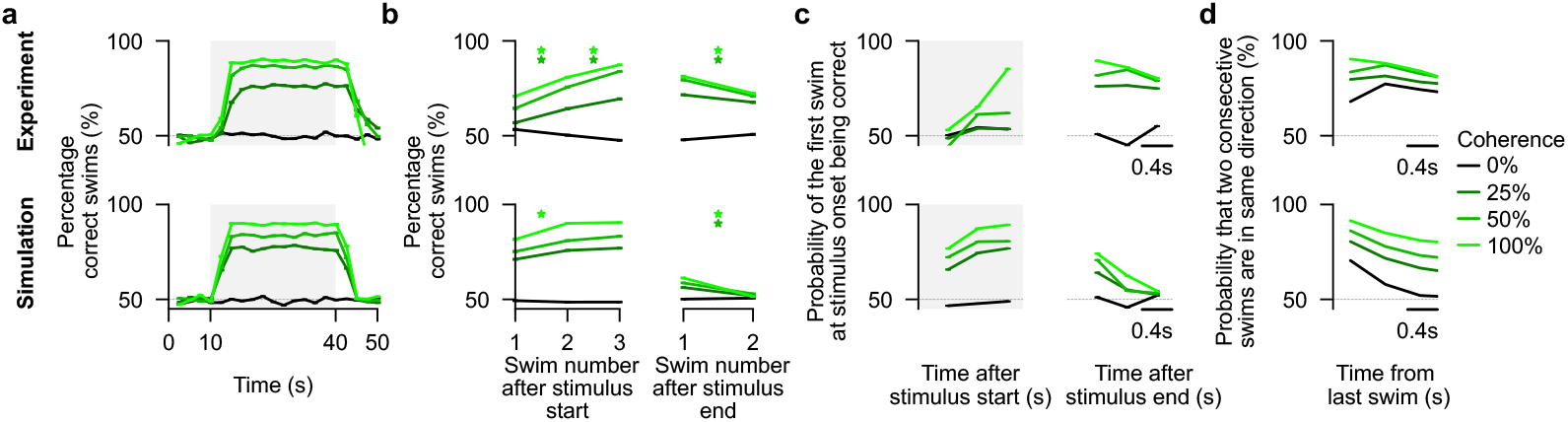
Fitted drift-diffusion models can predict behavioral features not explicitly used during model generation. **a**, Percentage of correct swims, binned over time (with bins spanning 2.5 s) for all tested coherence levels. Gray boxes indicate the period of motion coherence display. Before and after, we display 0% coherence. The slow rise and decay of the curve at the start and stop of stimulus presentation indicate temporal integration and persistent motion memory. **b**, Percentage of correct swims increasing in the first three swims after the start of the stimulus (left) and decreasing in the first two swims after the end of the stimulus (right). Asterisks (*) indicate significant (p < 0.05, bootstrapping hypothesis test) changes in performance from one swimming event to the next at a given coherence level (different green levels). **c**, Probability of the first swim after the stimulus start (left) and after the stimulus end (right) to be correct. Swimming events were binned depending on the delay relative to the stimulus start (left) or end (right). **d**, Probability to consecutively swim in the same direction as a function of inter-swim interval during periods of 0% coherence. The slow decay indicates the tendency to consecutively repeat the same sensorimotor decision. The top row in (a–d) represents results from experimental data, merged from all animals. The bottom row in (a–d) shows model-generated results, merged from individual model simulations. N=39 fish and N=39 individual fish-fitted models, respectively. All experimental animals were 5 dpf old. Same animals and optimized models as in related **Fig. 3**.

